# Environmental influences on the maximum quantum yield of terrestrial primary production

**DOI:** 10.1101/2023.11.11.566568

**Authors:** David Sandoval, Victor Flo, Catherine Morfopoulos, Iain Colin Prentice

**Affiliations:** Georgina Mace Centre for the Living Planet, Department of Life Sciences, Imperial College London, Ascot, UK; Department of Animal Biology, Plant Biology and Ecology, Autonomous University of Barcelona (UAB), E08193 Bellaterra (Cerdanyola del Vallès), Catalonia, Spain; CREAF, Ecological and Forestry Applications Research Centre, E08193 Bellaterra (Cerdanyola del Vallès), Catalonia, Spain; Department of Earth System Science, Tsinghua University, Beijing, China

## Abstract

**Summary:**

- Historically, terrestrial biosphere models (TBMs) have assigned the intrinsic (maximum) quantum yield of photosynthesis (φ_0_) a constant value for each plant functional type. However, experimental studies have shown that φ_0_– when measured on light-adapted leaves – depends on temperature. It is unclear whether this dependence is universal or biome-specific; how it is manifested at the ecosystem level; and how it should be represented in TBMs.
- By fitting empirical light-response curves to a global set of eddy-covariance CO_2_ flux measurements and correcting for photorespiration, we inferred apparent, ecosystem-level φ_0_values and their temperature responses across a wide range of environments.
- The temperature response of apparent ecosystem-level φ_0_ follows a universal bell-shaped curve. The shape of this curve does not markedly differ among biomes, but the maximum value of φ_0_ decreases with increasing aridity, its temperature optimum increases with increasing growth temperature, and its sensitivity to temperature increases as growth temperature declines.
- Our model for φ_0_(*T*) aligns with recent theory highlighting the role of cytochrome *b*_6_*f* in regulating the light reactions of photosynthesis. If implemented in TBMs, this model should allow better predictions of the responses of terrestrial ecosystem function to a warming climate.

## Introduction

Uncertainties in the prediction of photosynthesis can cause large discrepancies in projections of ecosystem responses to global environmental change (Harrison *et al*., 2021). Most current terrestrial biosphere models (TBMs) (Fisher *et al*., 2014; Harrison *et al*., 2021) rely on the biochemical theory of photosynthesis developed by Farquhar, von Caemmerer and Berry (FvCB) (Farquhar *et al*., 1980a). However, despite sharing the same core equations, ΤΒΜs have implemented them using varying conceptualizations of leaf-to-canopy scaling (from leaf-level photosynthesis to ecosystem-level gross primary production, GPP), and different assumptions about the values of key parameters that are not specified by the theory (Rogers *et al*., 2017). Hence, although TBMs based on FvCB might be expected to yield similar GPP estimates, in fact they do not. Indeed, many models show systematic differences both from one another and from accepted observational ranges, especially regarding the response of GPP to temperature (Dietze, 2014; Prentice *et al*., 2015; Harrison *et al*., 2021).

One cause of variation among models’ GPP estimates is the value or values assigned to a parameter of particular importance for the land carbon cycle (McNeall *et al*., 2024): the intrinsic quantum yield of photosynthetic carbon fixation (φ_0_). φ_0_ is the initial slope of the light-response curve of photosynthesis under non-photorespiratory conditions (high CO_2_ and low O_2_ partial pressure); thus, it sets the maximum efficiency of photosynthesis (Long *et al*., 1993; Rogers *et al*., 2017). We use the symbol φ_0_ here for the intrinsic, or maximum, quantum yield of CO_2_ assimilation at low light intensity to distinguish it from *φ* without a subscript, which denotes the quantum yield at low light intensity under ambient partial pressures of CO_2_ and O_2_. Both quantities are distinct from the maximum carboxylation rate (*V*_cmax_) and the maximum rate of electron transport (*J*_max_) as defined in the FvCB model, whose temperature dependencies have been studied far more extensively. Research on the light reactions of photosynthesis since the 1950s has established the theoretical maximum value for φ_0_to be either 1/8 = 0.125, i.e. one mole of assimilated CO_2_ requires at least eight moles of absorbed photons based on the NADPH requirement of carbon fixation (Warburg *et al*., 1950; Emerson, 1958; Walker, 1992; Hill & Govindjee, 2014), or 1/9 ≈ 0.111 based on the ATP requirement (Evans, 1987; Long *et al*., 1993). Almost all reported values have been obtained from measurements on leaves conducted in a non-photorespiratory atmosphere, within a narrow temperature range (25–30°C) and with adequate water supply. Under these conditions measured φ_0_ falls within the range of 0.07–0.125 mol CO_2_ mol^−1^ photon, with highest values in shade-adapted plants, and lower values in C_4_ compared to C_3_ plants (Ehleringer & Björkman, 1977; Ehleringer & Pearcy, 1983; Björkman & Demmig, 1987; Long *et al*., 1993; Singsaas *et al*., 2001; Skillman, 2008).

Despite limited knowledge of how φ_0_ responds to low or high temperatures and varying water availability, and how it upscales from leaves to ecosystems, it has generally been assumed that the available leaf-level observations are representative and sufficient to infer globally applicable values for modelling purposes (Walker, 1992; Long *et al*., 1993; Singsaas *et al*., 2001). This assumption has led to φ_0_ being prescribed in TBMs as a constant value per plant functional type, with no variation with temperature (Rogers *et al*., 2017). Empirical light use efficiency (LUE) models that incorporate a downward regulation factor for ambient CO_2_ (e.g. (Zhang *et al*., 2019); (Bao *et al*., 2022); (Cao *et al*., 2022) have similarly relied on prescribed, constant maximum LUE values for each plant functional type.

The temperature (in)dependence of φ_0_can be analysed using a corollary to the FvCB model (Farquhar *et al*., 1980b; von Caemmerer, 2000), which approximates the quantum yield of leaf-level photosynthesis at low light as follows:

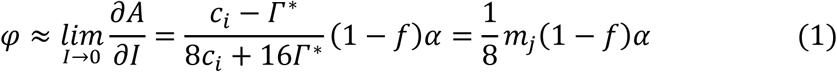

where *A* is the assimilation rate, *I* is the incident irradiance, *c_i_* is the intercellular CO_2_ partial pressure, α is the leaf absorptance, *f* is a correction factor for light quality (≈ 0.15: von Caemmerer, 2000) and *m_j_* = (*c_i_* − Г^∗^)⁄( *c_i_* + 2Г^∗^) where Г^∗^ is the photorespiratory compensation point, given by:

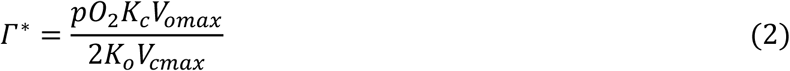

where *pO*_2_ is the partial pressure of oxygen, *V*__max_ and *K* are the maximum rates and Michaelis-Menten coefficients respectively, and the subscripts _o_ and _c_ refer to oxygenation or carboxylation. Under non-photorespiratory conditions Г^∗^ becomes negligible; so *m_j_* approaches 1.0, and φ_0_ approaches 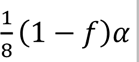 (Fig. 1a), independent of temperature. Under photorespiratory conditions φ is predicted only to decline with increasing temperature, due to the greater temperature sensitivity of *K*_o_ relative to *K*_c_.

**Fig. 1.**
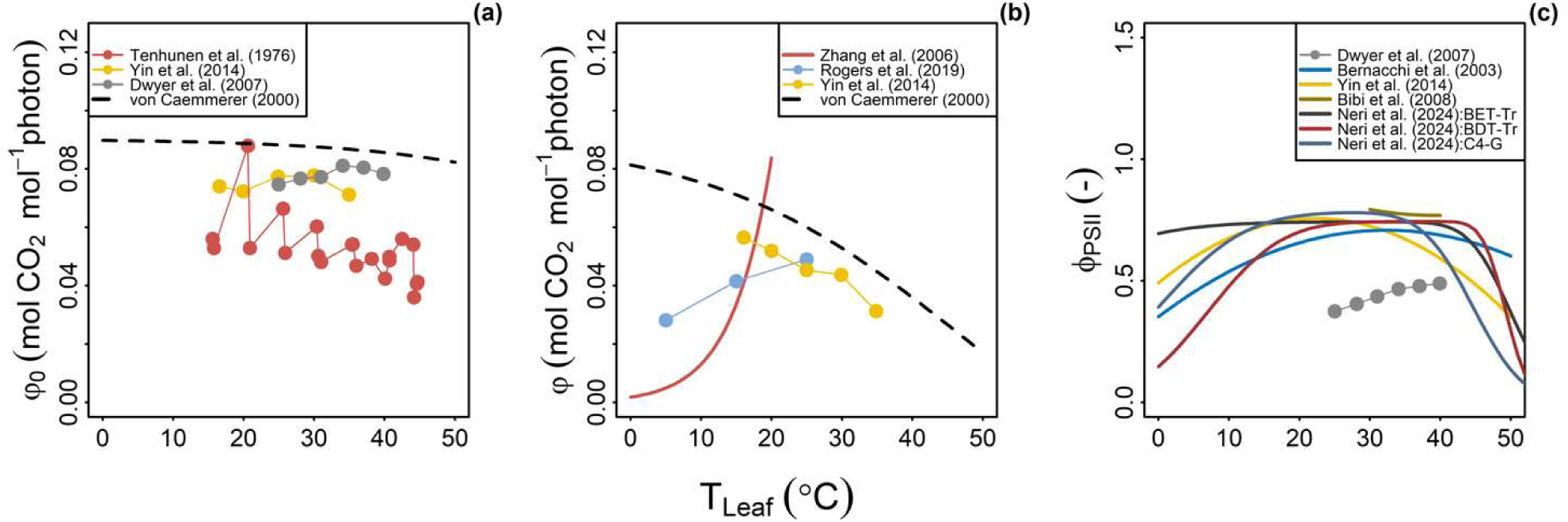
Maximum quantum yield responses to temperature from various sources. Connected dots represent observations, solid lines represent empirical models fitted to observations, and dashed lines represent predictions from the FvCB theory (Eq. (1)). **(a)** Non-photorespiratory maximum quantum yield of CO_2_ assimilation (Tenhunen *et al*., 1976b; von Caemmerer, 2000; Dwyer *et al*., 2007; Yin *et al*., 2014); Eq. (1) forced here with CO_2_ set at 750 μmol mol^−1^ and 2% O_2_. (**b)** Ambient maximum quantum yield of CO_2_ assimilation (von Caemmerer, 2000; Zhang *et al*., 2006; Yin *et al*., 2014; Rogers *et al*., 2019); Eq. (1) forced here with CO_2_ set at 400 μmol mol^−1^ and 21% O_2_. **(c)** Quantum yield of Photosystem II in light-adapted leaves from different studies (Bernacchi *et al*., 2003; Dwyer *et al*., 2007; Bibi *et al*., 2008; Yin *et al*., 2014; Neri *et al*., 2024). Selected PFTs from Neri *et al*. (2024): BET-Tr, broadleaf evergreen tropical tree; BDT-Tr, broadleaf deciduous tropical tree; C4-G, C4 grass.

However, these predictions are inconsistent with observations. Eq. (1) generally overestimates φ at low temperatures, under both non-photorespiratory and ambient CO_2_ conditions (Fig. 1b); and as noted by (Rogers *et al*., 2019) for Arctic plants, the observed change of φ with temperature below 25°C is opposite to that predicted, increasing rather than decreasing with temperature. This qualitative disagreement between theory and observations might explain some of the inconsistencies in the temperature responses of GPP predicted by FvCB-based TBMs (Bloomfield *et al*., 2023).

In principle, φ_0_reflects the emergent efficiency of the whole electron-transport chain. According to a recent mechanistic description of the electron transport system (Johnson & Berry, 2021), this involves the efficiencies of both Photosystems I and II, with the cytochrome *b*_6_*f* complex regulating the electron flow between the two in such a way as to minimise losses of absorbed light under low light while down-regulating electron transport (to match the use of ATP and NADPH for carboxylation) under saturating light. The different elements of the coupled photosynthetic system work in tandem and have distinct temperature dependencies (Johnson & Berry, 2021; Johnson *et al*., 2021). In the FvCB theory, by contrast, the only temperature effect applied to the electron-transport chain is to *J*_max_, which is empirically defined as the asymptotic rate of electron transport under saturating light and CO_2_ (Farquhar *et al*., 1980a; Johnson *et al*., 2021) – implying that φ_0_ is not or barely temperature-dependent, as suggested by Eq. (1).

Many empirical studies have focused on the initial stage of the electron-transport chain, the large protein complex Photosystem II, whose quantum efficiency (Φ_PSII_), defined as the number of electrons transported per photon absorbed by PSII (as measured by pulse amplitude modulation fluorometry) is expected to be proportional to φ_0_ when photosynthetic steady state is experimentally forced by actinic light under non-photorespiratory conditions (Björkman & Demmig, 1987; Genty *et al*., 1989; Oberhuber *et al*., 1993). When measured on dark-adapted leaves, Φ_PSII_shows little or no temperature dependence. In light-adapted leaves, however, Φ_PSII_ typically shows a distinct bell-shaped response to temperature (Fig. 1c) in both C_3_ and C_4_ species (Dwyer *et al*., 2007). Fig. 1 also shows data (or empirical models fitted to data) on φ_0_ (panel a), derived from CO_2_ assimilation measured under non-photorespiratory conditions, and φ (panel b), from CO_2_ assimilation under ambient conditions – indicating a range of responses to temperature but broadly suggestive of a unimodal pattern similar to that shown for Φ_PSII_ in panel c.

The large differences among the various published apparent quantum yield responses of *φ*_0_ and *φ* to temperature, the limited temperature range of many data sets at leaf-scale, and the disagreement between standard theory predictions and observations suggest that a more comprehensive global analysis would be helpful for the representation of φ_0_ in TBMs. Here we derive apparent, ecosystem-level values of φ, and thence φ_0_, from the slopes at low light of the light-response curves of net ecosystem exchange (NEE) as observed across the global network of eddy-covariance flux towers. We fit a universal empirical model to this response, and show how the model’s parameters are influenced by adaptation to different growth environments.

## Materials and methods

We exploited the abundance of half-hourly NEE data available from eddy-covariance CO_2_ flux measurements, together with *in situ* observations of the fraction of absorbed photosynthetic active radiation (fAPAR) made at 53 sites in the Ameriflux network (Novick *et al*., 2018). We estimated fAPAR from measured above- and below-canopy Photosynthetic Photon Flux Density (PPFD, μmol m^−2^ s^−1^) as follows (Carrara *et al*., 2018):

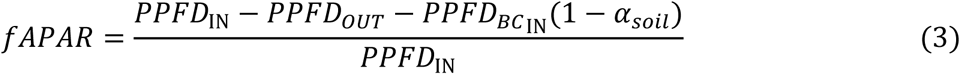

where PPFD_IN_ is the incoming (incident) PPFD and PPFD_OUT_ the outgoing PPFD at the top of the canopy, and PPFD_BCIN_ is the incoming PPFD below the canopy. α*_soil_* is the soil albedo in the visible light spectrum, taken here as 0.17 (Feister & Grewe, 1995). Absorbed PPFD (*I*_abs_) was estimated as the product of PPFD_IN_ and fAPAR. We additionally used a larger, globally distributed set of 310 sites (Pastorello *et al*., 2020; Hufkens, 2022; Ukkola *et al*., 2022) using remotely sensed fAPAR (Myneni *et al*., 2015) from the MODIS MCD15A3 product (Myneni *et al*., 2015), interpolated to daily timesteps.

For each site, we estimated φ_0_values within 1°C temperature bins using Nonlinear Least Squares. To capture the maximum efficiency and avoid limitations caused by factors other than temperature and light (e.g. soil moisture), we used only the upper envelope (95^th^ percentile) of the data; and only data points flagged as daytime good-quality observations were used. To account for the curvature of the light-response curve and ecosystem respiration (Re), we fitted the following hyperbolic equation to the NEE data (Smith, 1937; Stocker *et al*., 2020):

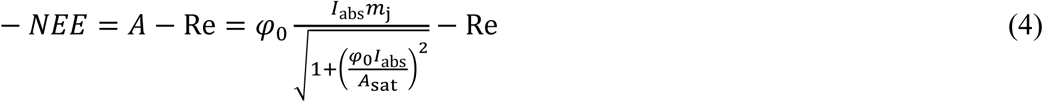

where *A*_sat_ is the asymptotic value of GPP under saturating light. The factor *m*_j_ corrects for photorespiratory conditions and the effect of vapour pressure deficit (*D*) on stomatal opening, following the least-cost hypothesis:

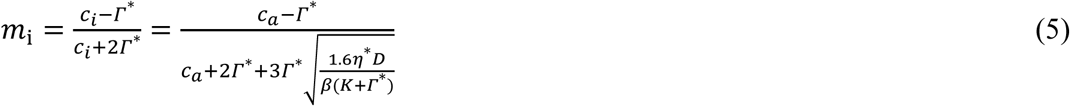

where *c*_a_ is the ambient partial pressure of CO_2_; *K* = *K*_c_ (1 + *p*O_2_/*K*_o_) is the effective Michaelis constant of Rubisco; η* is the viscosity of water, normalised by its value at 25°C; *D* is the vapour pressure deficit; and β is a constant, estimated as 146 (Stocker et al., 2020) based on global leaf δ^13^C data. The method is illustrated in Fig. 2.

**Fig. 2.**
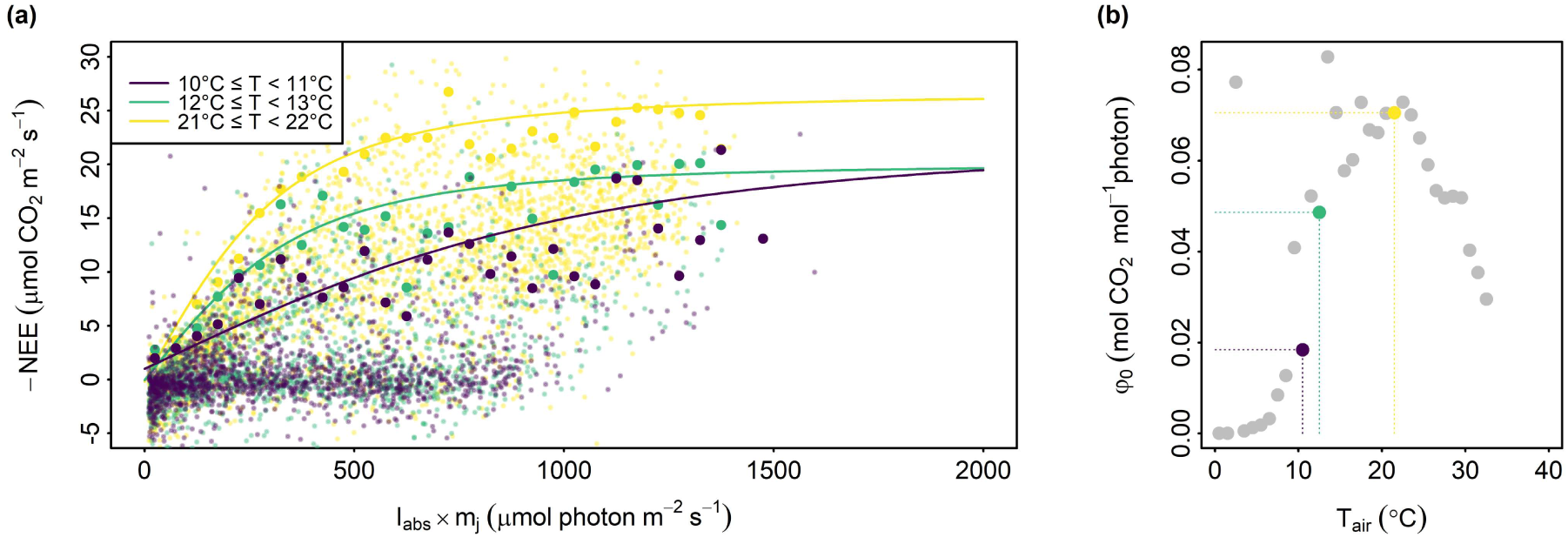
Method used to extract φ_0_(*T*). Three 1.0 °C intervals with different colours are shown as an example for the Harvard Forest site US-xHa (Biome: deciduous broadleaf forest, mean annual temperature = 6.6 °C, mean annual precipitation = 1071 mm). (**a)** Point clouds per temperature bin: larger points denote the 95^th^ percentile, and solid lines are Eq. (4) fitted to these 95^th^ percentiles. **(b)** φ_0_ extracted from the fitted Eq. (4) as shown in panel **(a)**.

Once φ_0_ was found at each temperature bin, we fitted φ_0_ = *f*(*T*) to a peaked Arrhenius equation (Eq. 6-7; Kattge & Knorr, 2007) by using the classic Nelder–Mead optimization algorithm (Nelder & Mead, 1965) with the Nash–Sutcliffe (NSE) coefficient (Nash & Sutcliffe, 1970), which relates the variance of the residuals to the variance of the data as the objective function, then we repeated this procedure for all sites:

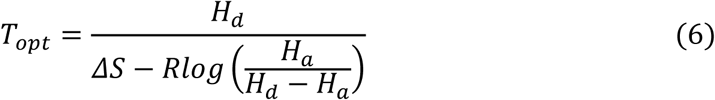

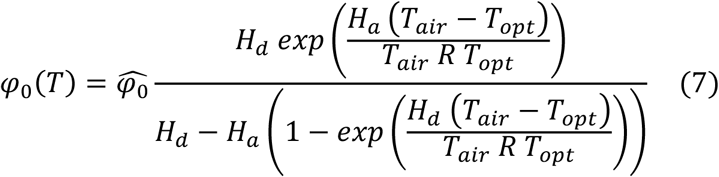

where *T_opt_* (K) is the temperature at the peak, *H_a_* (J mol^−1^) is the activation energy, *H_d_* (J mol^−1^) the deactivation energy, *ΔS* (J mol^−1^ K^−1^) is the entropy change between the ground and transition states of the reaction, φ̂_0_ is the intrinsic quantum yield at *T_opt_*, and *R* (J mol^−1^ K^−1^) is the universal gas constant.

To test whether the site’s bioclimate shapes this response to temperature, we fitted nonlinear models using bioclimatic indices explaining *ΔS*, *T_opt_* and φ̂_0_ parameters from the peaked Arrhenius equation with 10 000 bootstrap resampling iterations. The bioclimatic metrics tested were aridity index (*AI* = *PET*⁄*P*), evaporative index (*EI* = *ET*⁄*P*), evaporative fraction (*EI* = *ET*⁄*PET*) and mean temperature during growth days (mGDD_0_) where PET is the mean annual potential evapotranspiration, P is mean annual precipitation, and ET is the mean annual actual evapotranspiration. PET was estimated using the SPLASH v2.0 model (Sandoval *et al*., 2024), which accounts for surface temperature feedback on the net radiation. Actual evapotranspiration was estimated by converting the latent heat observed at the tower using the latent heat of vaporisation with corrections for local temperature and pressure. Growth days are defined as days with mean temperatures > 0°C. Mean growth temperature (mGDD_0_) was estimated as the average temperature during those days. All the environmental data used to estimate the bioclimatic metrics were measured *in situ,* except for sites where the records were shorter than five years. In such cases, data from ERA5 (Muñoz-Sabater, 2019) were used instead.

To compare our model, theoretical simulations of φ_0_ were carried out using the model and code described in (Johnson *et al*., 2021) with default parameters for *Populus fremontii* as used in (Johnson & Berry, 2021). Bioclimatic metrics for the sites described in (Johnson & Berry, 2021) and (Rogers *et al*., 2019) were calculated using data from ERA5 (Muñoz-Sabater, 2019) and SPLASH v2.0 (Sandoval *et al*., 2024). To evaluate the effect of the new formulation on global GPP φ_0_(*T*) was implemented in the P model and global simulations were run with both the original which applies Bernacchi *et al*. (2003) φ_0_(*T*) correction, and the new formulation.

## Results

φ_0_ varied considerably across biomes and climate zones. Mean values showed a smooth transition between biomes, with higher values in forests and lower values in grasslands and shrublands (Fig. 3a). Examining this transition using climate zones, a marked pattern emerged, driven by the aridity gradient. Tropical rainforest (Af) and tropical monsoon (Am) climates appeared at the upper end of the φ_0_ axis, followed by sites with “no dry season” (*f* types), and then by sites with dry/wet seasonality (*w* and *s* types). Finally, sites from arid climates (BW*) and steppe appeared in the lower end (BS*) (Fig. 3b and 3c).

**Fig. 3.**
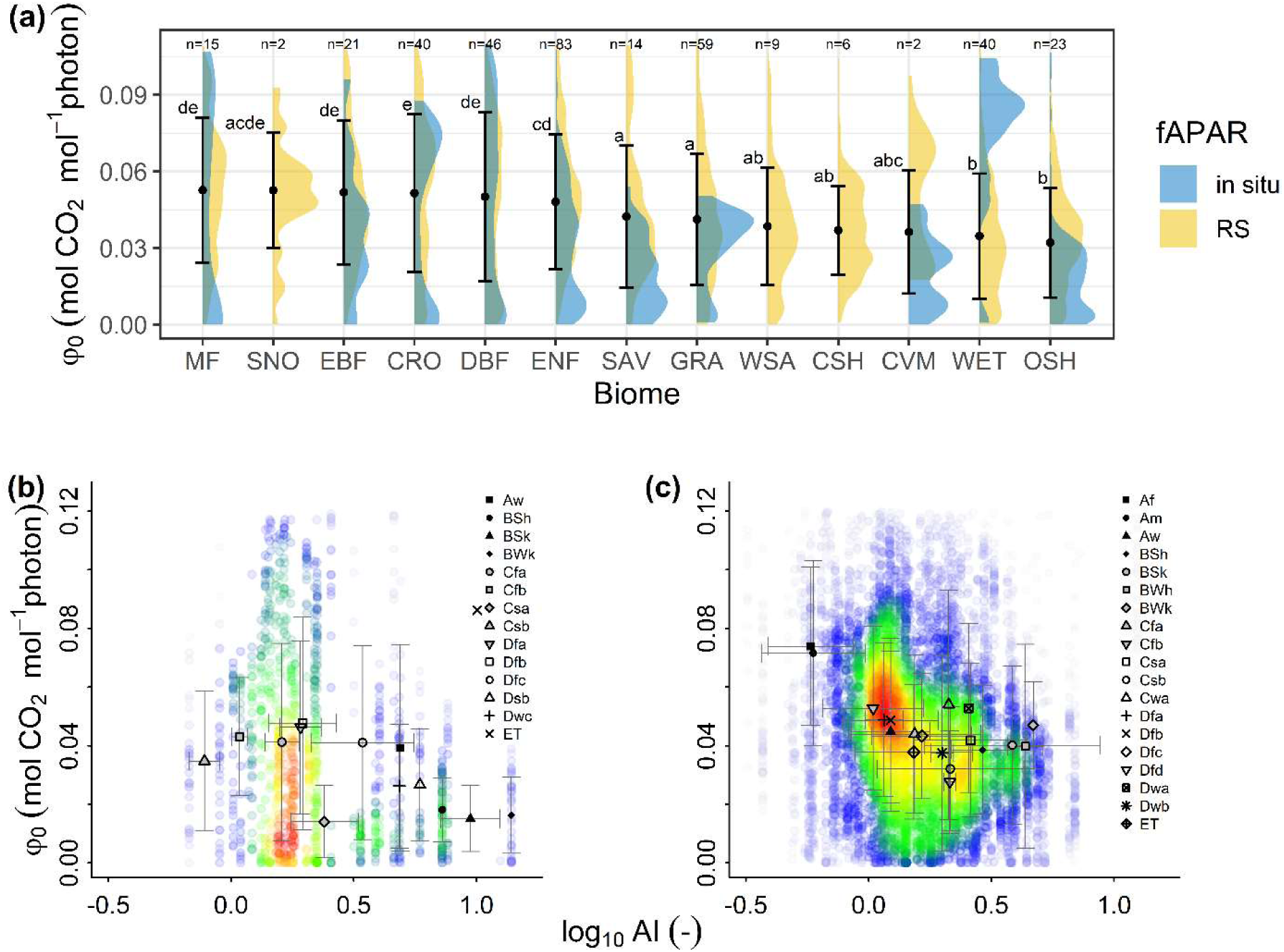
Observed φ_0_ inferred by using *in situ* and remotely sensed fAPAR. **(a)** Probability distribution of φ_0_ per biome and fAPAR source. The symbols and the black bars denote the mean and standard deviation per biome, based on combined fAPAR sources. Post hoc Tukey test results at the top of the bars and the number of sites at the top of the distributions. (**b)** φ_0_as a function of aridity index from in situ fAPAR sites. **(c)** φ_0_ as a function of aridity index from RS fAPAR sites. Symbols in panels **b** and **c** show the average φ_0_values per climate zone and the colours denote point density.

The response of φ_0_to temperature showed a bell-shaped pattern across all sites (Fig. 4a). In general, the average φ_0_ steadily increased up to a maximum, followed by a sharper drop. This pattern also emerged using the 75^th^ and 95^th^ percentiles of φ_0_(Fig. 4a). The global maximum efficiency, defined based on the 95^th^ percentile, was 0.111 ± 0.003 mol CO_2_ mol^−1^ photon and was located at the optimal temperature of 20.46 ± 0.37 °C. The bell-shaped response remained consistent across biomes with the in situ fAPAR data, while with the RS fAPAR wetlands and savannas showed inconsistencies (Fig 4b). Open shrublands (OSH) showed lower φ_0_values and lower sensitivity to *T*_air_, whereas mixed forests (MF) showed the highest φ_0_ values and higher sensitivity to *T*_air_.

**Fig. 4.**
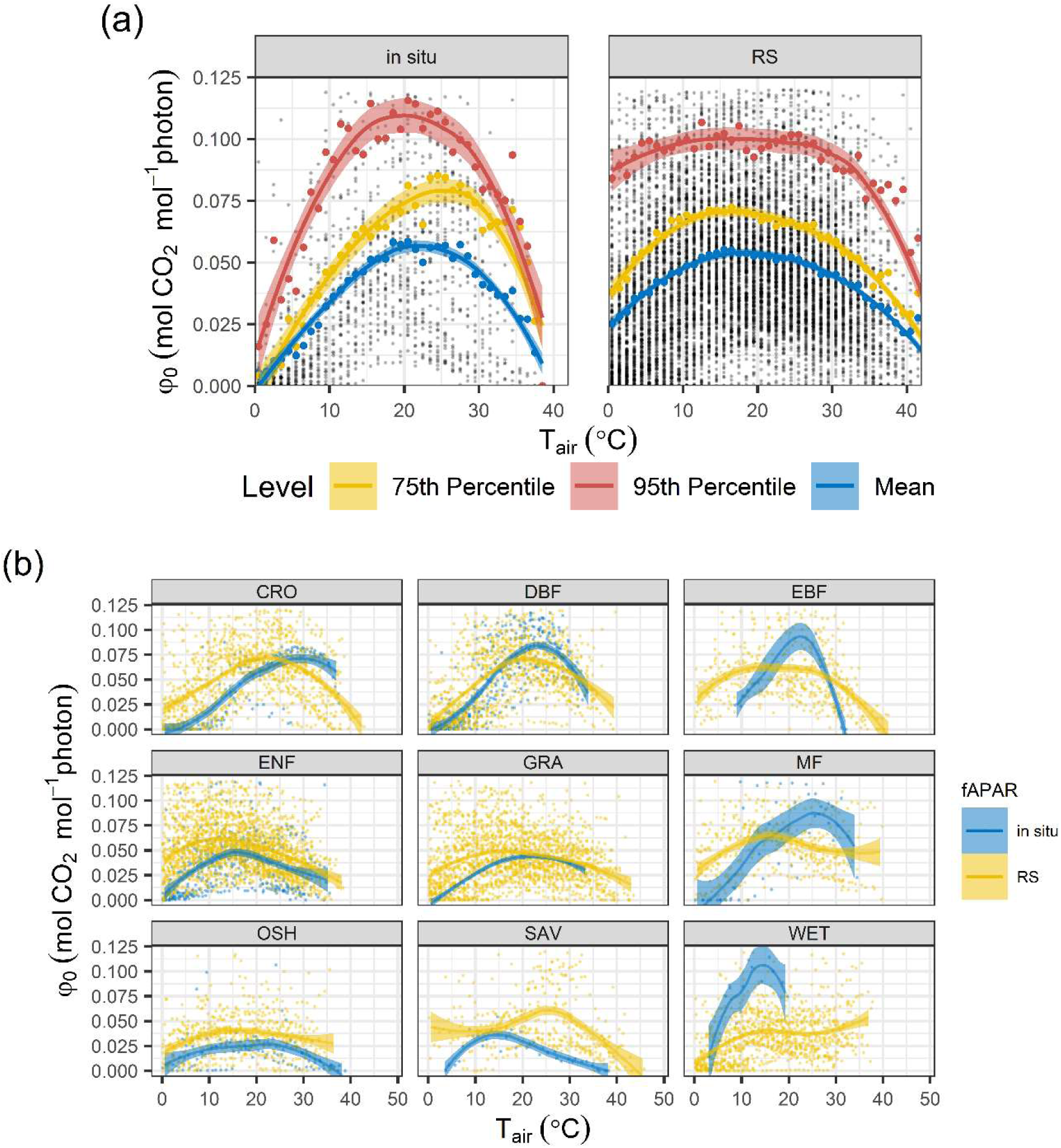
φ_0_ responses to temperature using all data pooled. The ribbons show a LOESS smoothing function including its 95% confidence intervals. **(a)** Pooled observations from sites with *in situ* fAPAR and sites with remotely sensed fAPAR. **(b)** Pooled observations per biome and fAPAR source.

These patterns are less pronounced when the (temporally sparse) remote-sensing fAPAR data are used. In ENF, GRA and EBF biomes, remotely sensed fAPAR produces higher φ_0_ at low temperatures – possibly indicating multiple optimal temperatures within each biome. In biomes where about the same temperature range was covered by both datasets (DBF, MF and OSH), the temperature sensitivity of φ_0_(*T*) was also similar between the datasets.

We further analysed the temperature response at individual sites using the fitted peaked Arrhenius curves. The shapes of the response were consistent regardless of the location, biome or fAPAR source. However, different temperature sensitivities and optimal temperatures emerged. For example, wetlands (WET) as depicted in Figure 5 show the extreme difference between a wetland in the Peruvian rain forest (PE-QFR) and a wetland in Greenland (GL-Zaf).

**Fig. 5.**
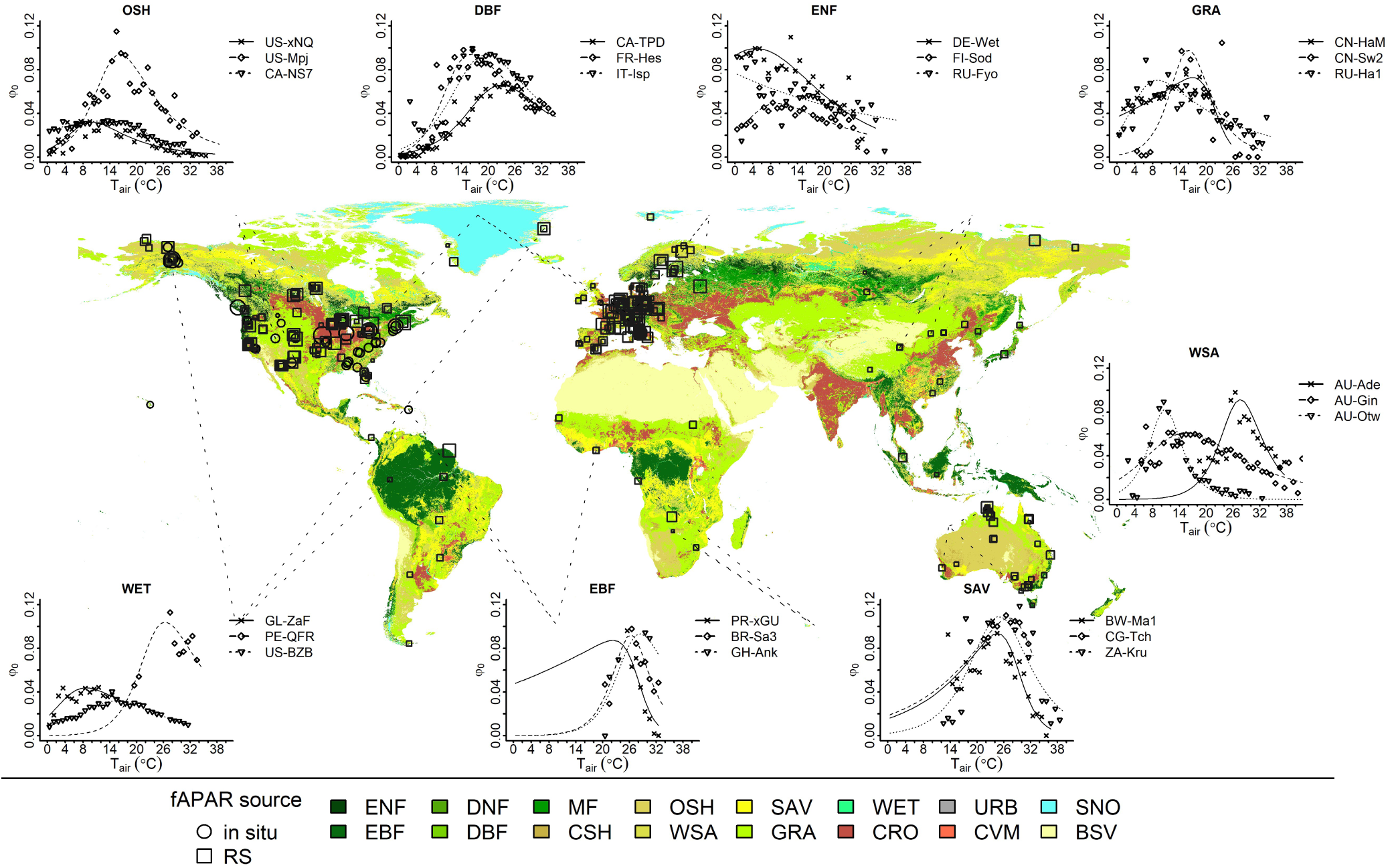
φ_0_ responses to temperature using peaked-Arrhenius curves. Observation (points) and predictions (lines) from selected sites.

Globally, the curves fitted site by site to the peaked Arrhenius equation were able to explain φ_0_(*T*) with reasonably good accuracy. Performance was slightly better for the *in situ* fAPAR, explaining 85% of the variation while explaining 72% of the variation for the *RS* fAPAR. Most of the mean values (observed and simulated) for biomes fell close to the 1:1 line, except for OSH which was slightly overestimated and WET underestimated (Fig. 6a).

**Fig. 6.**
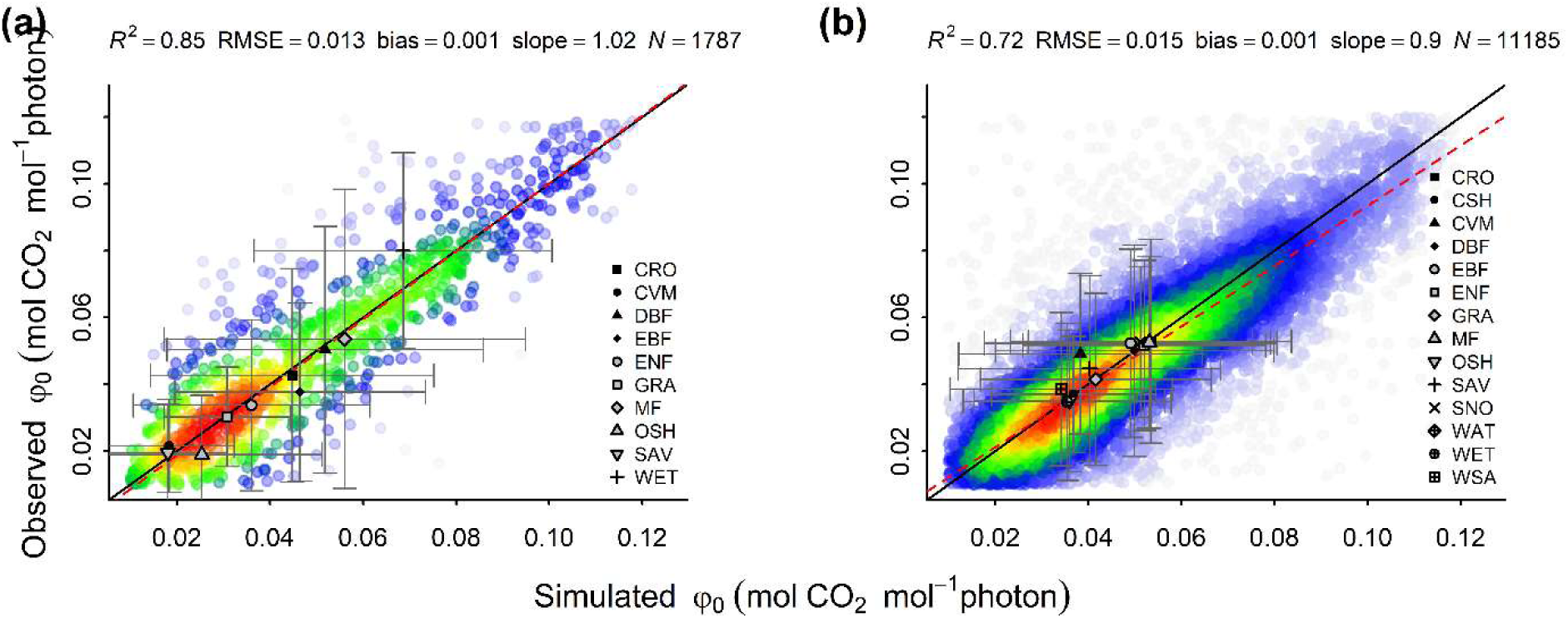
Correlations of observed and simulated φ_0_(*T*) using peaked Arrhenius curves. **(a)** using *in situ* fAPAR observations. **(b)** using remotely sensed fAPAR observations. Colours denote point density.

The peak φ_0_ (φ͡_0_) showed considerable variation among sites, while *H_a_* was relatively stable among sites. For most (350 sites) *H_a_* was ∼ 70.9 kJ mol^−1^. *H_d_* varied from ∼ 111 kJ mol^−1^ to 351 kJ mol^−1^, while Δ*S* varied from ∼ 0.2 kJ mol^-1^ K^-1^ to >1.5 kJ mol^-1^ K^-1^ and was highly correlated to *H_d_*. φ̂_0_ decreased from a plateau when AI ≤ 1 with increasing aridity following an inverse sigmoid response (Eq. 8):

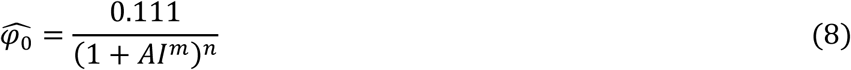

where *m* = 4.09 (95% CI: 4.07 - 9.99) and *n* = 0.12 (95% CI: 0.050 - 0.169). Thus φ̂_0_ shows a steep decay from 0.111 when AI > 1, stabilizing (when AI > 8) at a value around 0.03 (Fig. 7a). Some of the sites that fall outside the 95% confidence intervals are tagged in the plots and discussed in the next section. The global average of φ̂_0_, according to the prediction from equation (12) (Fig. 7b), was 0.08 mol CO_2_ mol^−1^ photon.

**Fig. 7.**
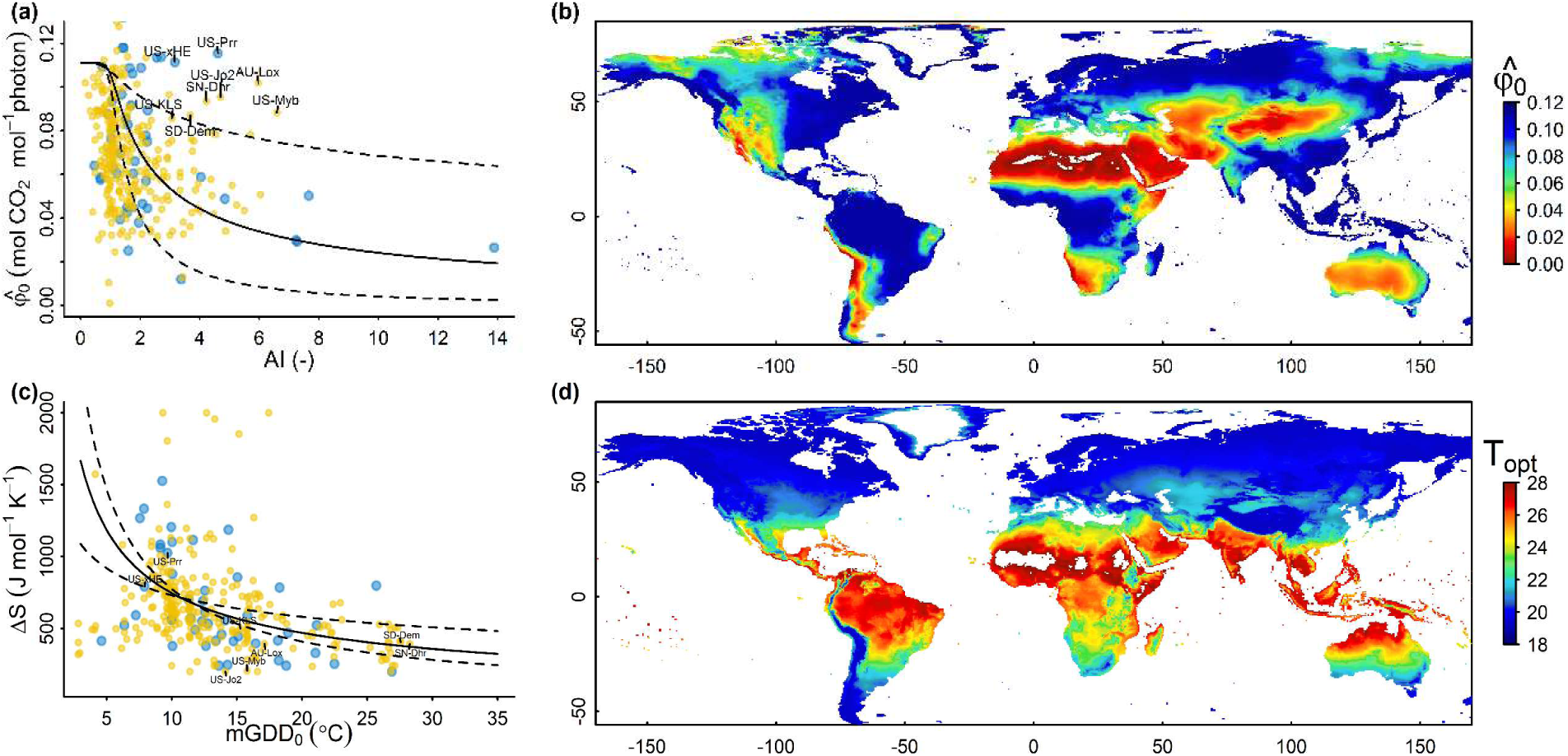
Bioclimatic patterns of the peaked Arrhenius parameters. The colours in the scatter plots refers to fAPAR measured *in situ* or from remote sensing. **(a)** φ̂_0_ response to aridity: solid line regression, dashed lines 95% confidence intervals. **(b)** Global prediction of φ̂_0_ using equation (8). **(c)** Entropy change (Δ*S*) variation with growth temperature: solid line regression, dashed lines 95% confidence intervals. **(d)** Global prediction of optimum temperature for φ_0_using equation (9).

*ΔS* decreases non-linearly, approaching stability for mGDD_0_ > 30 °C, as described by Eq. (9), where *ΔS*_0_ = 3468.2 (95% CI: 1523.9 – 6408.2) and kS = −0.668 (95% CI: 0.321 - 0.639):

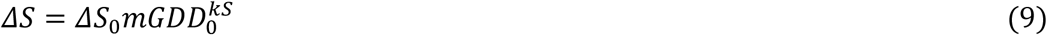

Thus, replacing Eq. (9) in Eq. (6) produces *T_opt_* within a range from ∼17 °C in boreal regions to ∼26 °C in the tropics (Fig. 7d), which also results in a rate of change 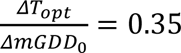.

## Discussion

We derived apparent φ_0_at the ecosystem level for globally distributed sites, using half-hourly *in situ* measurements of fAPAR and eddy-covariance measurements of CO_2_ exchange. We found that it shows a bell-shaped response with temperature, and this pattern remains consistent across all individual sites analysed, even when remotely sensed fAPAR was used in the absence of *in situ* fAPAR measurements (Fig. 5). However, this pattern was less consistent when the data were grouped per biome (Fig. 4b), suggesting that the temperature response curve for φ_0_ is not biome-specific but instead gradually modulated by bioclimate. In particular, increasing aridity reduces φ̂_0_, while higher growth temperatures shift the optimum temperature upward and simultaneously decrease the sensitivity of φ_0_to instantaneous temperature. The use of absorbed rather than incident radiation should implicitly account for the effect of high diffuse fractions during low sun angles and their confounding effect with low temperatures. This effect should be clearer with the *in situ* fAPAR observations, which are higher under greater diffuse light, independently of temperature. Since MODIS overpasses occur at 10:30 and 13:30, it is unlikely that they captured early-morning periods of high diffuse light and their effect on fAPAR. However, this remains a caveat, as no data were available on the partitioning between diffuse and direct radiation.

No previous studies have reported φ_0_at ecosystem scale. However in our analysis, the fitted maximum value of φ_0_ (0.111 mol CO_2_ mol^−1^ photon) is identical to the theoretical maximum limited by ATP generation of 0.111 (Long *et al*., 1993) and to the maximum observed by (Walker, 1992). It also closely matches other values reported for single leaves under unstressed conditions including (Björkman & Demmig, 1987) (0.106 on average for C3 plants and 0.112 for conifers), (Skillman, 2008) (0.106) and (Singsaas *et al*., 2001) (0.108).

Our analyses show that φ_0_is generally higher in forests, followed by grasslands and shrublands, suggesting that φ_0_ varies among species from different life forms – in line with the observations by (Singsaas *et al*., 2001), who noted that even a 100% variation in photorespiration plus alternative electron sinks could not fully explain the observed variation in φ_0_ among species. This result contrasts with the findings of (Björkman & Demmig, 1987), (Walker, 1992) and (Long *et al*., 1993) who suggest an almost constant φ_0_ among all species, regardless of their life-form. However, since our estimations were done at ecosystem scale, some of this difference could also reflect distinct canopy structures: a closed canopy would prevent leaves in the lower levels from being exposed to excessive light that could cause photodamage, thereby increasing the overall ecosystem-level φ_0_ (Long *et al*., 1994; Dashti *et al*., 2025). Another source of variation could be attributed to different proportions of C3/C4 plants which have distinct φ_0_ (Oberhuber *et al*., 1993), with higher proportions of C4, eq. (5) might overestimate φ_0_ in tropical grasslands and savannas due to C4 plants’ low photorespiration.

Our findings of a universal peaked response to temperature are consistent with the predictions from the theory developed by (Johnson & Berry, 2021) (Fig 8c) – supporting the idea that the bell-shaped response of φ_0_ to temperature is a general plant response which propagates to the ecosystem, thus creating a general ecosystem response to temperature.

**Fig. 8.**
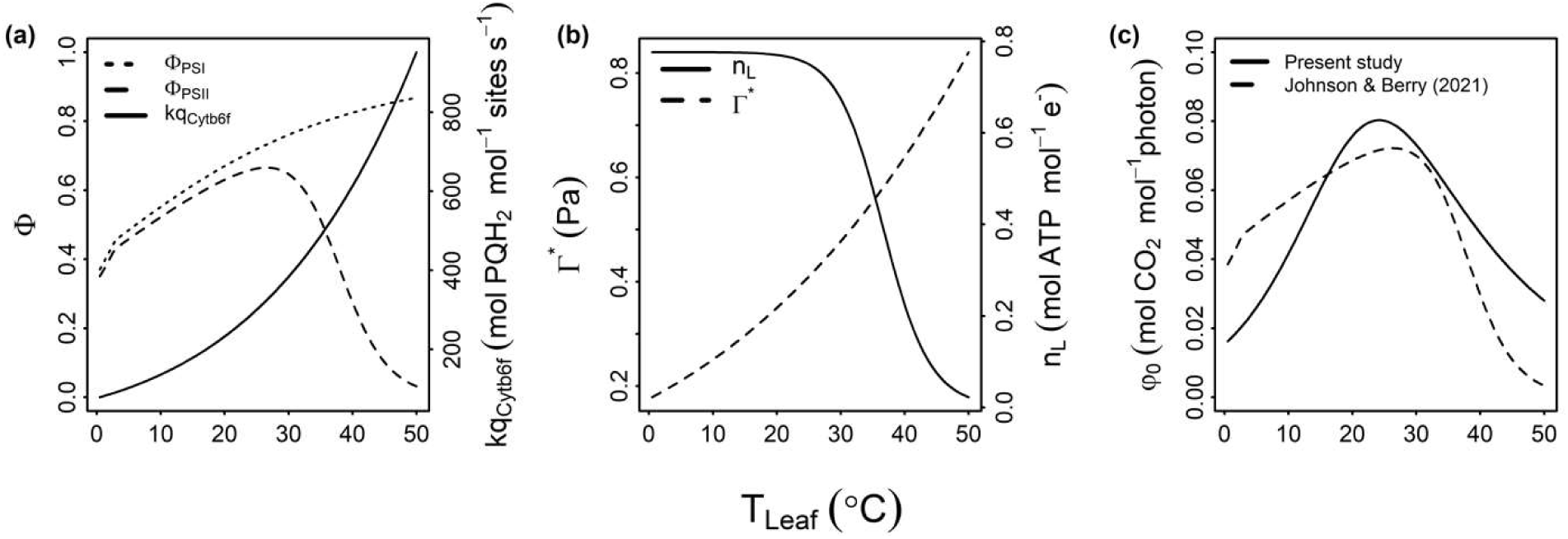
Theoretical responses of selected electron-transport chain components to temperature, following Johnson & Berry (2021). Simulations were carried out under non-photorespiratory conditions: CO_2_ = 700 μmol mol^−1^, *pO_2_* = 2029 Pa, and low light (150 μmol m^−2^ s^−1^). **(a)** Efficiencies of photosystems I and II and the catalytic constant for cytochrome *b*_6_*f*. **(b)** The photorespiratory compensation point and the coupling efficiency of linear electron flow to the proton circuit (ɳL). **(c)** φ_0_(*T*) predictions from theory and from the present study.

In the Johnson-Berry theory, cytochrome *b*_6_*f* (Cyt *b*_6_*f*) modulates the transfer of electrons from plastoquinol (PQH2) to oxidized plastocyanin (PCox) at a rate *kq* (Fig. 8a), redistributing excitation energy between photosystems and diverting energy to thermal dissipation. Cyt *b*_6_*f* also couples this electron transfer to the proton circuit from the stroma into the lumen, modulating the coupling between linear and cyclic electron flux (Johnson & Berry, 2021). When light is limiting, Cyt *b*_6_*f* activity is at its maximum and so *kq* depends primarily on temperature, following a regular Arrhenius response. Most of the electron flow passing through Cyt *b*_6_*f*, at limiting light, and connecting to the proton circuit, is thought to be linear; this efficiency decays exponentially at high temperatures (Fig. 8b) (Johnson & Berry, 2021).

Low φ_0_ at low temperature could be explained by the accumulation of sustained non-photochemical quenching (NPQ) (Porcar-Castell, 2011) or by the rate of repair of photosystem II, which slows down at low temperature (Greer *et al*., 1986; Mattila *et al*., 2020). If this rate of repair becomes slower than the rate of PSII damage, it could lead to photoinhibition even under low light (Osmond, 1994; Tyystjärvi & Aro, 1996). This damage-repair imbalance is also documented when the rate of damage is enhanced by high temperature (Murata *et al*., 2007; Sage & Kubien, 2007). It has been reported that PSI activity decreases at low temperatures only in cold-sensitive plants (Sonoike, 1999), otherwise it is thermally stable. Its inactivation does not occur before the complete inactivation of PSII (Berry & Bjorkman, 1980; Oberhuber & Edwards, 1993).

Although we corrected for stomatal closure and photorespiration, implicitly accounting for Rubisco deactivation at high temperature, other factors could affect φ_0_(*T*) such as the solubility of CO_2_ in water: this decreases as temperature increases, reducing the CO_2_ concentration in C_3_ plants and favouring oxygenation by Rubisco, as noted by (Ehleringer & Björkman, 1977). However, this effect is counterbalanced by the increase in CO_2_ diffusivity and hence the increase of mesophyll conductance (von Caemmerer & Evans, 2015).

According to Eq. (6), respiration offsets the light response curve in each temperature bin without affecting the slope φ_0_. Nonetheless, we double-checked our results by repeating the analysis using GPP from the FLUXNET dataset and setting respiration to zero, producing very similar results (Fig. S4).

Our results indicate that the maximum quantum efficiency *φ̂*0 at the ecosystem level decreases as climatological aridity increases, consistent with the observations of (Fu *et al*., 2022) regarding the maximum evaporative fraction, and with (Mengoli *et al*., 2023) regarding the maximum light-use efficiency of GPP. Several explanations are possible, including: (a) persistent effects of drought on the photosynthetic apparatus, such as damage to the light-harvesting complexes of PSII and reduction in the size of the PSI antenna (Hu *et al*., 2023); (b) high leaf mass per area (LMA) (Wright *et al*., 2004), typical in plants adapted to aridity, favouring photoprotective thermal dissipation over photochemistry (Adams *et al*., 1987; Valladares *et al*., 2000); (c) coordination of hydraulic and allocation traits (Flo *et al*., 2021) such that plants of dry environments minimize hydraulic risk (Bassiouni *et al*., 2023).

A few sites, however, showed relatively high φ̂_0_ at AI > 1. Although we cannot account for every case, there are many possible reasons why local conditions might differ from the bioclimate of the surrounding regions. For example, some sites such as AU-Lox (Stevens *et al*., 2011) and US-KLS (Ji *et al*., 2021) are irrigated; while others, such as SN-Dhr (Tagesson *et al*., 2015) and US-Jo2 (Pérez-Ruiz *et al*., 2022), experience a strong seasonal concentration of precipitation. Some sites have access to groundwater, like SD-Dem (Ardo *et al*., 2008), or are man-made wetlands, like US-Myb (Eichelmann *et al*., 2018). Other sites show a high PET/P ratio, but a significant portion of the energy is used for melting snow and thawing permafrost such as US-xHE (Osterkamp *et al*., 2009) and US-Prr (Nakai *et al*., 2013).

Activation energies for φ_0_ showed little variation among sites; this is also the case for the activation energies of *V*_cmax_ and *J*_max_, as reported by (Kattge & Knorr, 2007). Δ*S* decreases with increasing growth temperature, as also reported for *V*_cmax_ and *J*_max_ by (Kattge & Knorr, 2007) and (Kumarathunge *et al*., 2019). Variation in *ΔS* shifts the optimum temperature by 0.35 °C per degree of growth temperature and simultaneously reduces the sensitivity of φ_0_ to T, as also noted by (Yin & Struik, 2009). This response is consistent with the observations of increased thermal stability of PSII at higher growth temperatures due to changes in thylakoid membrane lipids (Berry & Bjorkman, 1980). It is also consistent with the effect of higher growth temperatures on the accumulation of zeaxanthin, which enhances nonphotochemical quenching and thermal energy dissipation in PSII (Demmig-Adams *et al*., 1989; Busch *et al*., 2007). Overall, the variations of *ΔS* and *φ͡*0 with mGDD_0_ and aridity respectively support our general hypothesis that the temperature response of φ_0_(*T*) depends on adaptation to bioclimate.

We replicated our analysis on a year-by-year basis in sites with at least ten years of *in situ* fAPAR records (*n* = 3) (Fig. S1) and did not observe any signal of acclimation. While only three sites may not provide sufficient evidence, this observation is consistent with what was reported by (Bernacchi *et al*., 2003). It would imply that the acclimation of *V*_cmax_ to temperature, which is a well-documented phenomenon (Medlyn *et al*., 2002; Kattge & Knorr, 2007), does not apply to φ_0_. However, some studies have reported acclimation to temperature of the initial slope of the light-response curve, such as (Dwyer *et al*., 2007) and (Herrmann *et al*., 2020). To our knowledge, only (Tenhunen *et al*., 1976a) fitted the same equation as in this study and found variation of φ_0_ with temperature – but the model presented there assumed constant φ_0_for simplicity.

Comparing our predictions with independent observations by (Rogers *et al*., 2019) (Fig. 9) shows that considering the temperature dependency of φ_0_eliminates a general problem shared by TBMs, of overpredicting φ_0_at low temperatures. Our model also predicts a sharper decline in φ_0_ at temperatures higher than *T*_opt_. All else equal, this translates into a sharper decrease in assimilation at high temperatures.

**Fig. 9.**
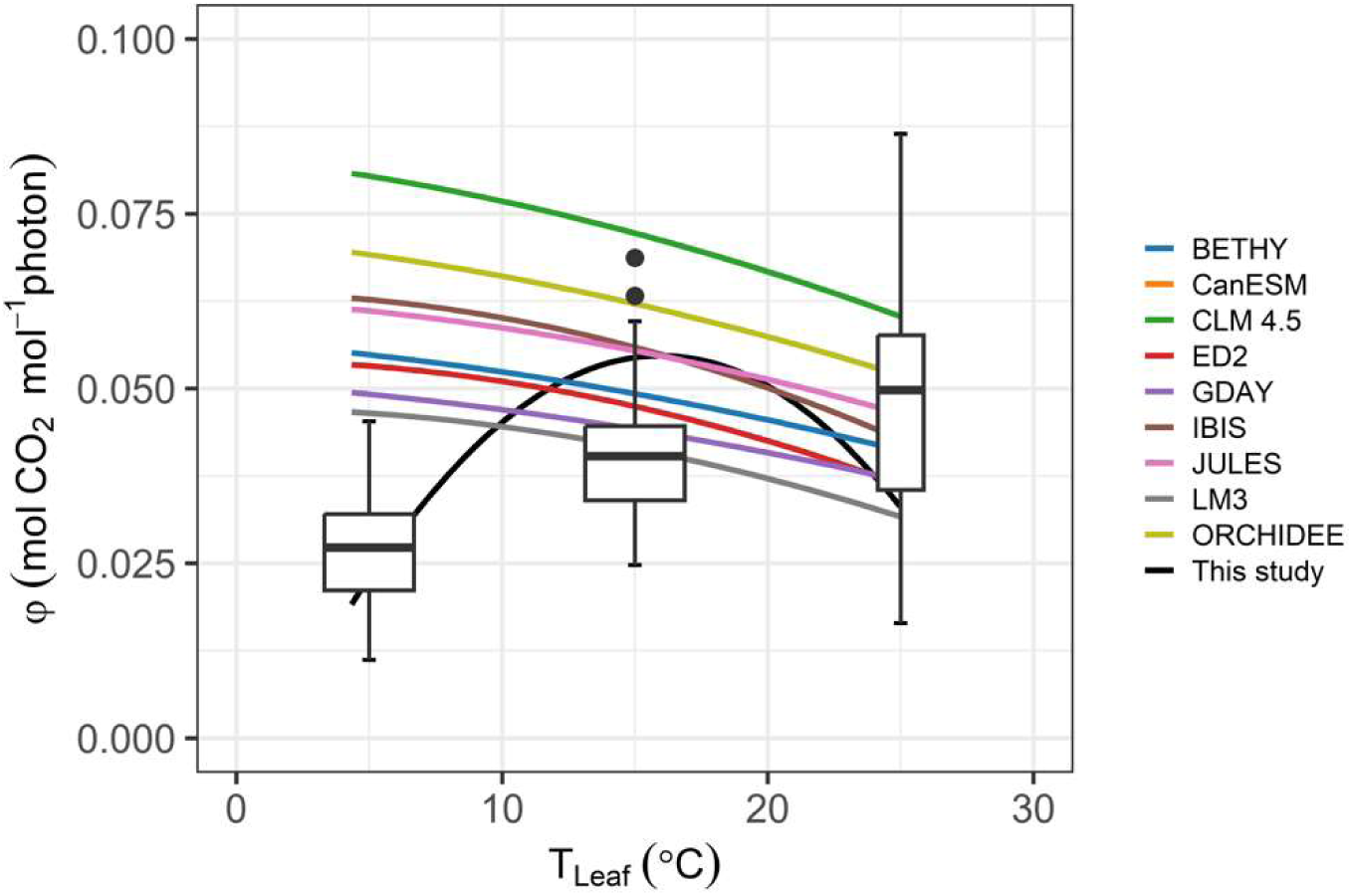
Realized quantum efficiency of primary production as predicted by different models. The boxes and whiskers show measurements from (Rogers *et al*., 2019) on Arctic plants; the black line shows results of our model. Results of other TBMs (coloured lines) are based on parameters and settings from (Rogers *et al*., 2019).

When this new formulation, with global coefficients (Eqs. 6 to 9), was tested on the P-model without any soil moisture limitations, we simulated an increase in global total annual GPP from 114.3 ± 3.49 PgC yr^-1^ to 161.0 ± 4.38 PgC yr^-1^, with respect to the original. The new value is closer to values inferred from ^14^C (∼160 PgC yr^-1^), ^18^O (150–175 PgC yr^-1^) and carbonyl sulfide uptake, COS (157 ± 8.5 PgC yr^-1^), but far from values derived from inverted soil respiration (149 ± 29 PgC yr^-1^), and inferred from SIF (135.5 ± 8.8 PgC yr^-1^) (Li & Xiao, 2019; Graven *et al*., 2024; Lai *et al*., 2024) (Fig. 10a). This change, however, was unevenly distributed; the increase was apparent in the tropical rainforests where the annual GPP increased by 1–3 kgC m^-2^ yr^-1^, while in temperate and boreal forests the increment was moderated to ∼0.5 kgC m^-2^ yr^-1^. In arid regions, in contrast, there was a moderate decrease in modelled GPP (Fig 10b).

**Fig. 10.**
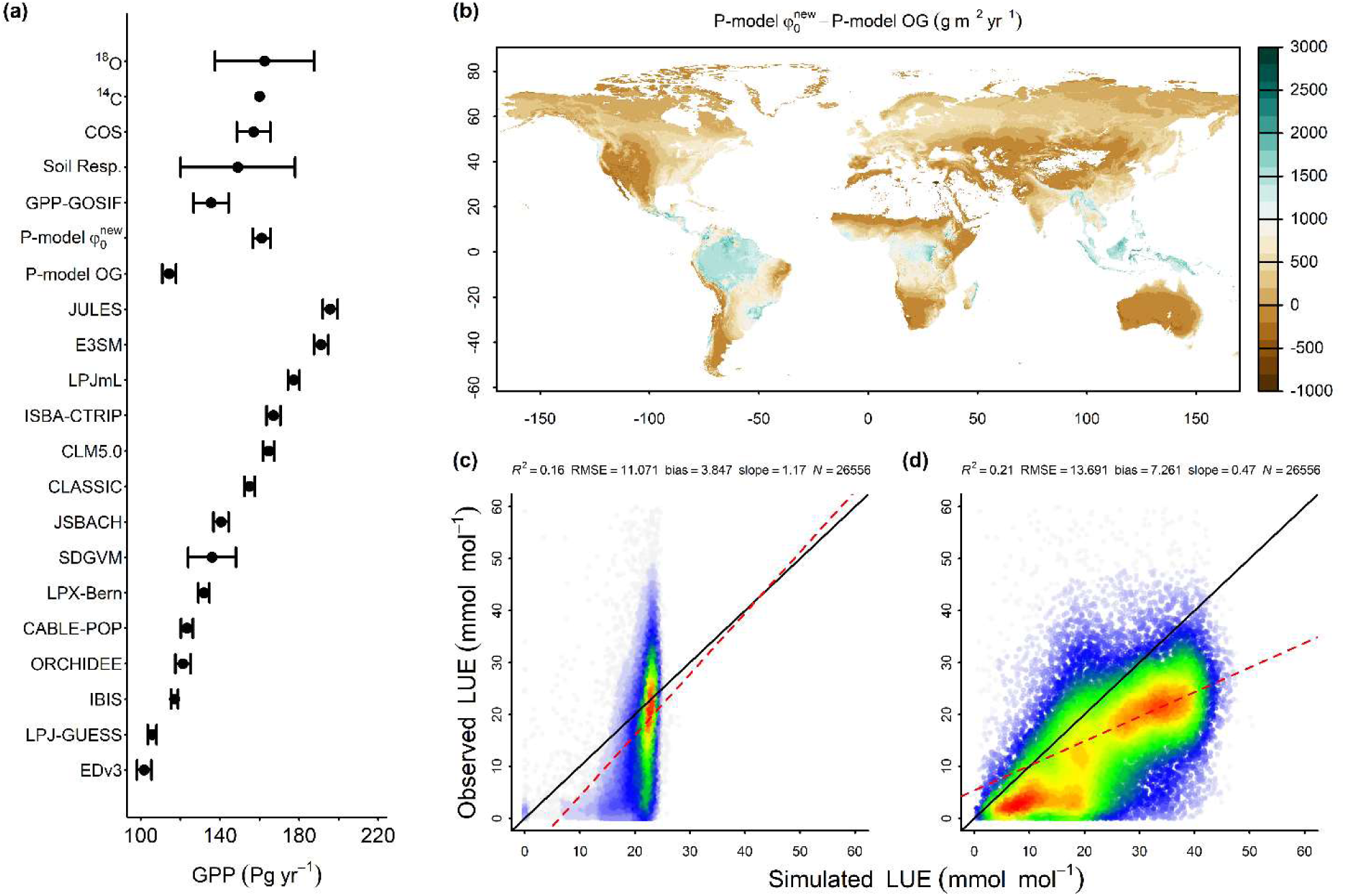
φ_0_implementation in the P model. **(a)** Global mean annual GPP from different sources, including the TRENDY models (Sitch *et al*., 2024) and this study. **(b)** Spatial distribution of the difference between global mean annual GPP, averaged over the years 2003 to 2020, from the new and the original P model (φ͡_0_ = 0.087 and *f* (T) from Bernacchi *et al*., 2003). **(c)** Correlation of observed and simulated LUE monthly values using all the flux-tower data pooled, original P-model φ_0_. **(d)** Correlation of observed and simulated LUE monthly values using all the flux tower data pooled and the new P model φ_0_.

When we compared monthly LUE from the P-model with the original vs. the new φ_0_ formulation, we saw an improvement in the *R*^2^ from 0.16 to 0.21 (Fig 10d). However, the slope and the bias were degraded compared to the original. A possible reason is the fAPAR source. For example, (Stocker *et al*., 2020) found a significant change in the magnitude of GPP when switching from MODIS fAPAR to fAPAR 3g. In our results, the underestimation occurs for high LUE values, which often represent tropical rainforests – where the error in remotely sensed fAPAR due to constant cloudiness is well documented (Zhang-Zheng *et al*., 2024). It is worth noting that in (Stocker *et al*., 2020) the same evaluation was done, but only roughly one-third of the data was available at that time; their R^2^ was 0.31. In the same way, in our evaluation the original formulation was not able to surpass ∼25 mmol mol^-1^ of monthly LUE (Fig 10c).

Overall, we have provided global evidence that φ_0_ is not a fixed parameter (or fixed per plant functional type) as has been widely assumed in TBMs. Instead, it has a bell-shaped response to temperature, whose parameters are determined by bioclimatic adaptation. Incorporating these responses will be key to reliably assessing the response of the terrestrial biosphere to continuing climate change.

## Acknowledgments

This work is a contribution to the LEMONTREE (Land Ecosystem Models based On New Theory, obseRvations and ExperimEnts) project. This research received support through Schmidt Sciences, LLC (Grant number G-21-61881: DSC, ICP, CM).

## Author contributions

Conceptualisation, coding and first draft writing: DS. All authors provided scientific input, reviewed and edited the final manuscript.

## Data and materials availability

All data, is freely available at The code is available at Supplementary Material

