## Supplementary Material for "Environmental influences on the maximum quantum yield of terrestrial primary production"

Sandoval, D. *et al.*

**This PDF file includes:**

Figs. S1 to S4

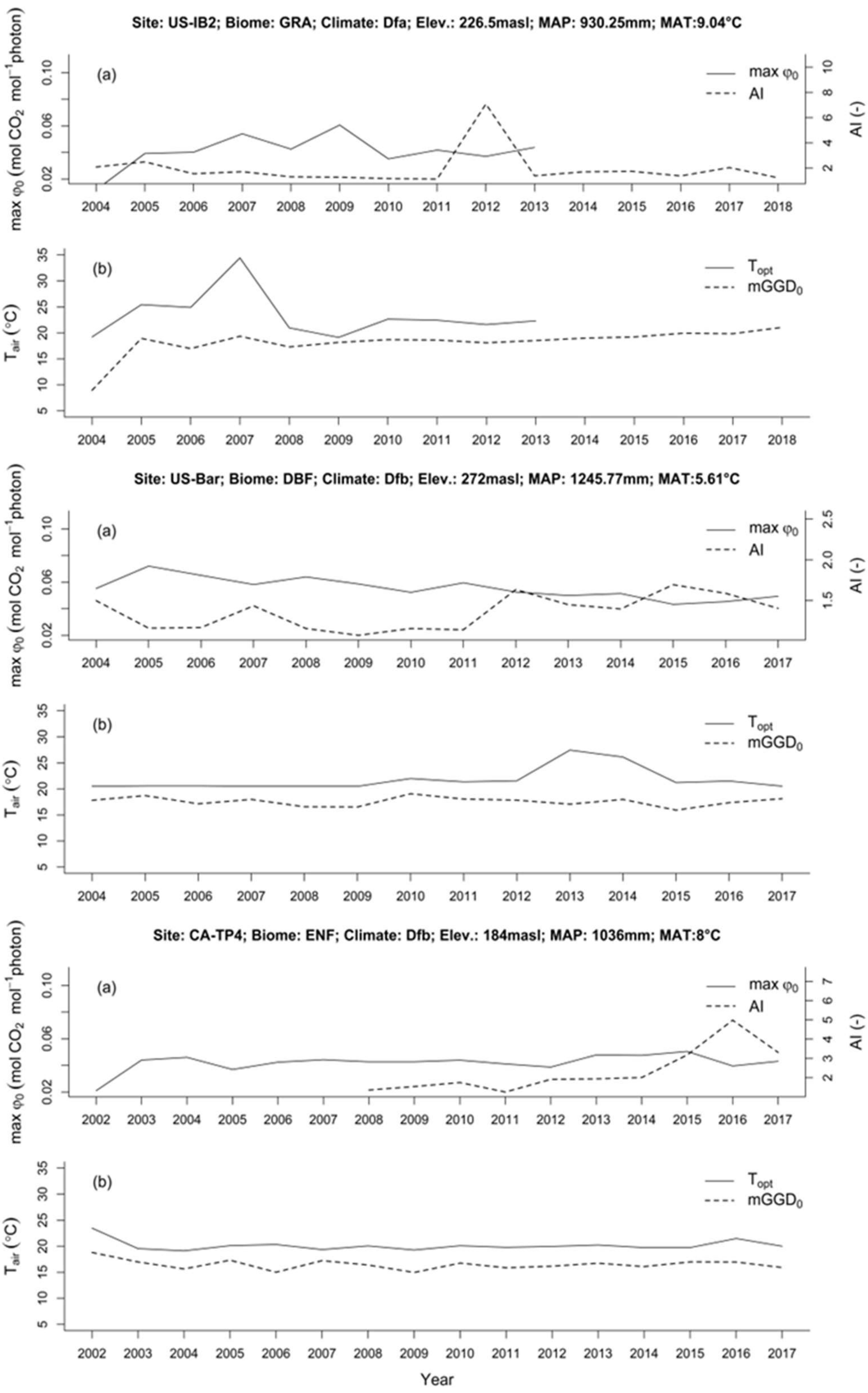


**Fig. S1.** Temporal dynamics of the parameters shaping $\varphi_0(T)$ at a selected sites

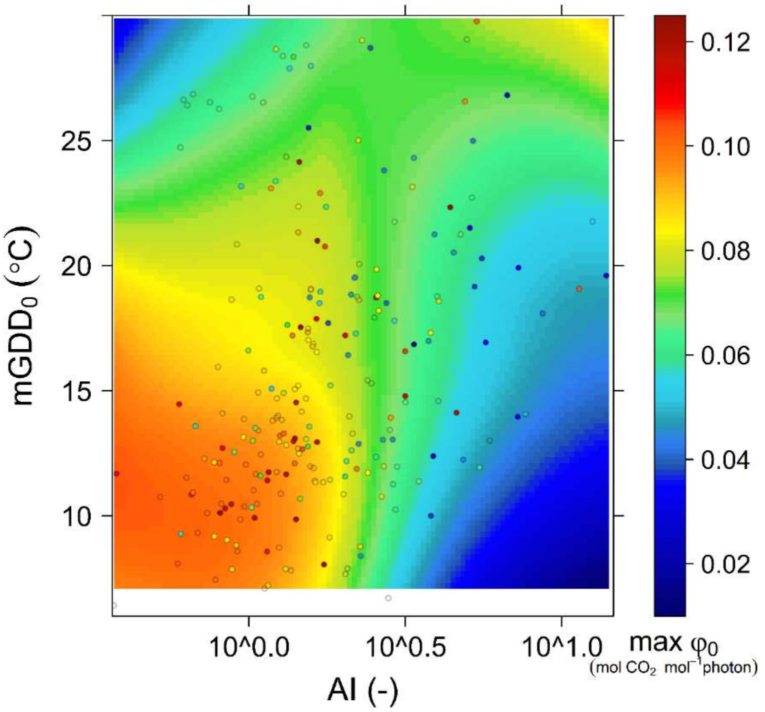


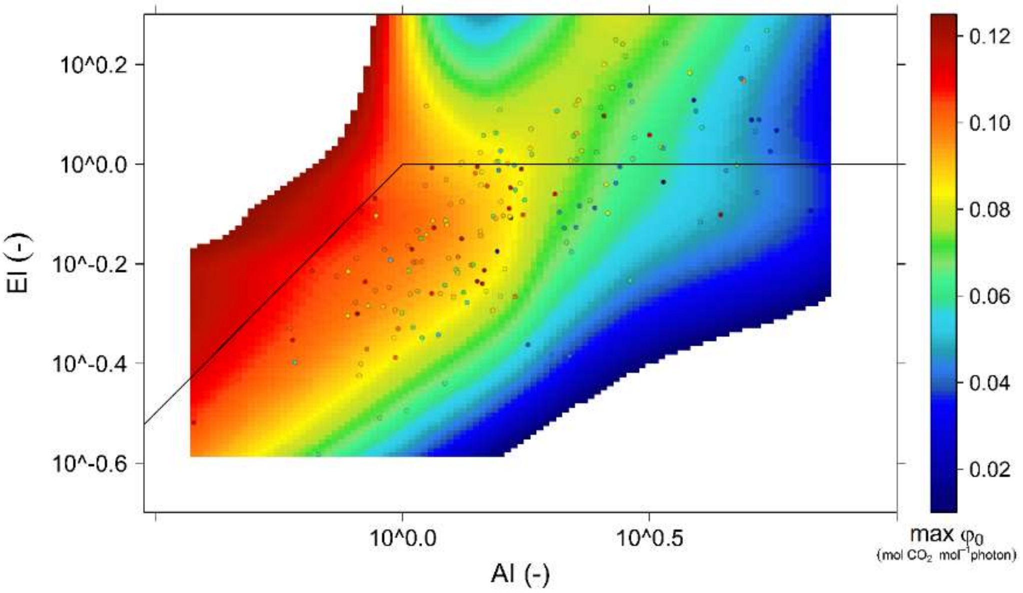


**Fig S2.** $\widehat{\varphi_0}$ depicted in climate spaces. (a) $\widehat{\varphi_0}$ in the space aridity-growth temperature. (b) $\widehat{\varphi_0}$ in the Budyko space

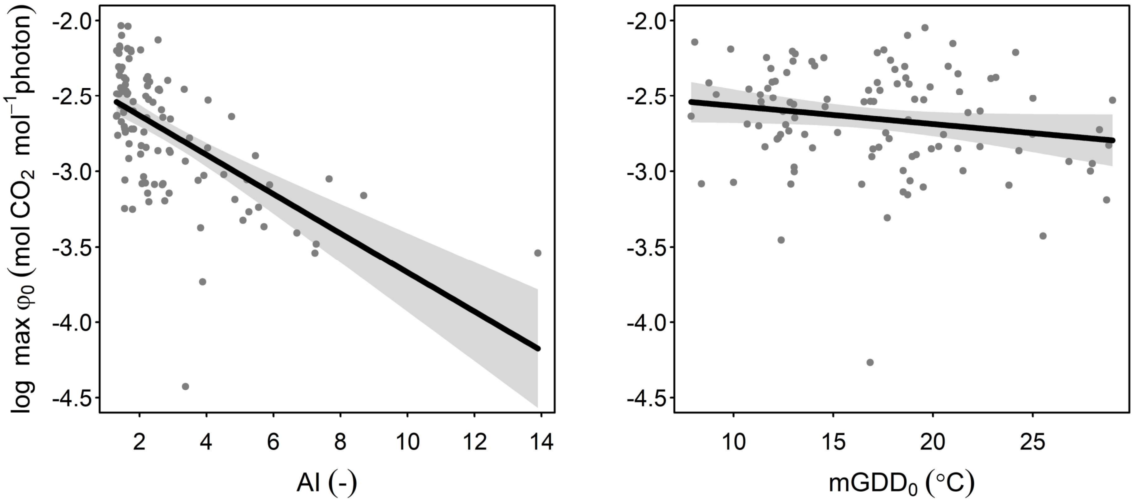


**Fig. S3.** Other patterns in the PA parameters. **a** partial residuals plot of $\widehat{\varphi_0}$ and AI. **b** partial residuals plot of $\widehat{\varphi_0}$ and $mGDD_0$.

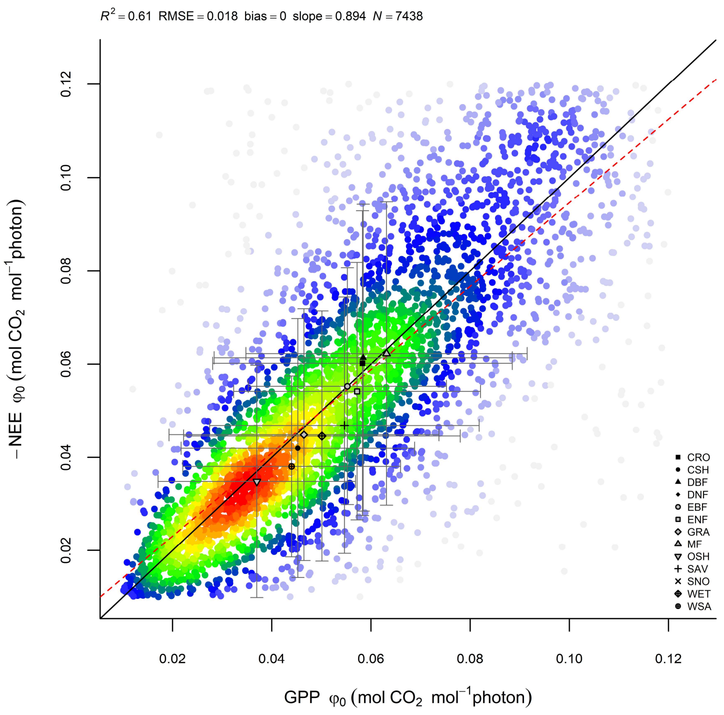


**Fig. S4.** Correlation of estimated $\varphi_0$ with NEE and estimated with GPP using the FLUXNET dataset.
